# Increased homozygosity due to endogamy results in fitness consequences in a human population

**DOI:** 10.1101/2022.07.25.501261

**Authors:** N.A. Swinford, S.P. Prall, C.M. Williams, J. Sheehama, B.A. Scelza, B. M. Henn

## Abstract

Recessive alleles have been shown to directly affect both human Mendelian disease phenotypes and complex traits like height. Pedigree studies also suggest that consanguinity results in increased childhood mortality and adverse health phenotypes, presumably through penetrance of recessive mutations. Here, we test whether the accumulation of homozygous, recessive alleles decreases reproductive success in a human population. We address this question among the Namibian Himba, an endogamous agro-pastoralist population, who until very recently practiced natural fertility. Using a sample of 681 individuals, we show that Himba exhibit elevated levels of “inbreeding”, calculated as the fraction of the genome in runs of homozygosity (F_ROH_). Many individuals contain multiple long segments of ROH in their genomes, indicating that their parents had high kinship coefficients. However, we did not find evidence that this is explained by first-cousin consanguinity, despite a reported social preference for cross-cousin marriages. Rather, we show that elevated haplotype sharing in the Himba is due to a bottleneck, likely in the past 60 generations. We test whether increased recessive mutation load results in observed fitness consequences by assessing the effects of F_ROH_ on completed fertility in a cohort of post-reproductive women (n=69). We find that higher F_ROH_ is significantly associated with lower fertility among women who have had at least one child (p<0.006). Our data suggest a multi-locus genetic effect on fitness driven by the expression of deleterious recessive alleles, especially those in long ROH. However, these effects are not the result of consanguinity but rather elevated background identity by descent.

## Introduction

Through the process of mutation, deleterious alleles are constantly arising in a population, and an understanding of how these variants affect phenotype is crucial for the study of human evolution and genetic architecture. Extensive previous work has been done to understand the variance in mutation load among populations, the demographic models that drive those loads, the distribution of fitness effects for new mutations, and the efficacy of selection for removing deleterious variants. However, direct evidence of the consequences of mutation load on fitness in human populations is lacking (1–6). Genetic theory dictates many factors affect a population’s mutation load, including founder effects, bottlenecks, small population size, endogamy, and consanguinity. Because a population’s mutation load relates to the number of deleterious mutations that have accumulated in its gene pool over time (1), it is thought that an increased mutation load leads to fitness consequences. Further, many deleterious mutations tend to be recessive (2) so processes that increase the level of homozygosity in a genome may result in strong fitness consequences. Evidence for the deleterious effects of recessive mutation load has been shown in multiple species. Increased homozygosity is linked to higher mortality in translocated desert tortoises (7) and decreased fitness in Florida scrub-jays (8). Decreased fitness has also been observed in *Drosophila* in mutation accumulation experiments (9). In humans, increased homozygosity has been related to decreased height and cardiometabolic disease phenotypes (10, 11). Additionally, it is well known that higher incidences of recessive Mendelian disorders are typically found in founder or endogamous populations (12–14). “Inbreeding” refers to mating between individuals who share one or more common ancestors, and among human populations it is frequently used to describe consanguineous unions (those between individuals related up to the degree of 2^nd^ cousins) or populations which have experienced recent founder effects (14–16). Here, we reserved the term inbreeding to refer to either the formal process of ‘inbreeding depression’ or close familial unions.

The reduction in fitness associated with an individual’s mutation load is dependent on the level of dominance of individual alleles. Previous work has shown that estimates of mutation load can greatly differ under additive versus recessive models (3, 6). Furthermore, there is a relationship between a mutation’s level of dominance and deleteriousness—the more deleterious a mutation, the more recessive it tends to be. Work in various non-human species has also shown that mildly deleterious alleles are often at least partially recessive, with an average dominance level of *h*=0.25 (2). Szpiech et al (17) looked at human populations and demonstrated that long runs of homozygosity (ROH) contained disproportionately more deleterious variants than short runs. Additionally, it has been estimated that all human individuals carry between at least 3 and 5 alleles that, if made homozygous, would be lethal (4). Because consanguinity or founder effects increases the likelihood that an individual will inherit identical segments from both parents, resulting in long ROH, inbred individuals would be expected to carry a higher burden of recessive deleterious, and possibly lethal, genotypes. Thus, these processes highlight the impact of recessive mutation load in a population. Levels of inbreeding depression can therefore be calculated as the fraction of the genome in ROH (F_ROH_) (12, 18).

It is important to note, however, that evidence currently exists for both biological costs and benefits associated with inbreeding depression. These costs include negatively affecting fitness in many species by decreasing offspring viability or reproductive success (8, 14, 19–22) and can exhibit differential intensity based on sex. For example, in Florida scrub-jays and an isolated human population in the Swiss Alps, females were more strongly affected (8, 22). Specifically, prior human studies using pedigree data have shown that increased parental relatedness, such as via consanguinity, increases childhood mortality and can decrease the total number of surviving children (16, 21, 22). However, Helgason et al showed that 3^rd^-4^th^ degree cousins have higher fitness than either more closely or more distantly related individuals (i.e. a “Goldilocks” zone) as assessed from Icelandic pedigrees (23). It is unclear from the Icelandic pedigrees whether this reflects biological benefit or whether cultural factors, such as the proximity of nearby kin, increased reproductive success.

Other studies drawn from cross-cultural anthropology, have demonstrated that there can be important cultural benefits associated with consanguinity, which is common both historically and contemporarily. In order to more fully understand the consequences of inbreeding, the biological effects on fitness must be considered alongside potential social and economic benefits of marriage between kin. Consanguineous marriage, typically among cross-cousins, is currently found among more than 10% of the world’s population (24) but occurs at much higher rates in parts of Africa and Asia (25, 26). Consanguineous marriage has been shown to provide many social and economic benefits, including the ability to maintain power and resources within families, and as a strategy for risk reduction (19, 21, 25, 27–29). Societies with more consanguineous marriage tend to have denser, more intensive kinship networks, which help prevent the dilution of material wealth and promote resource-based defense (26). Some studies report higher fertility for consanguineous couples, but these findings may be explained by other factors such as lower age at marriage and first birth (25).

To assess whether recessive mutation load results in observable fitness consequences in a human population, we combined demographic and genetic data (n = 681) from a community of Himba pastoralists, an endogamous, semi-nomadic population residing in northwestern Namibia. The Himba are expected to be previously bottlenecked and practice polygyny with a reported first-cousin preference for arranged marriages, but “love matches” exist as well (30). They exhibit an unusually high rate of extra-pair paternity (31), and half-sibling (and other second degree) relationships constitute a very large proportion of all the relationships among individuals (32), increasing the risk that reproduction could occur between these relatives. Additionally, until recently, they were a natural fertility population, and continue to have a pronatalist ideology and fairly short interbirth intervals (between 1 and 3 years) (33).

Our study differs from prior work on fertility by a) utilizing genome-wide data rather than pedigree-based inference of inbreeding coefficients, b) assessing the effect of recessive deleterious mutations in unions which span a range of kinship coefficients rather than consanguineous unions, c) incorporating data from a natural fertility population (i.e. no hormonal contraceptive use), and d) modeling the background levels of genetic drift in the population.

## Results

We find elevated levels of homozygosity in the Himba as measured by F_ROH_ for three different minimum length thresholds: 500 kb, 1500 kb, and 5000 kb (Figure 1). The lengths of runs of homozygosity (ROH) segments reflect the time since a common ancestor, where shorter ROH result from events occurring in the more distant past and longer ROH result from more recent events. In an idealized outbred population, the F_ROH_ expectation for the offspring of first cousins would be 0.0625, and F_ROH_ calculated using a minimum threshold of 1500 kb to call ROH is comparable to a pedigree estimate of inbreeding (13). However, this measure is not applicable to a population like the Himba who have elevated levels of IBD sharing (see below). Therefore, to estimate the F_ROH_ expectation for the offspring of Himba first cousins, we added 0.0625 to the average amount of background relatedness in the population resulting from more distant demographic history. To calculate this background level of inbreeding, we set a low (500 kb) threshold to call ROH and calculated F_ROH_ for all individuals. We then filtered out the individuals with F_ROH_ levels greater than 0.0625 and calculated the average level of F_ROH_ in the remaining individuals (0.0420). This background F_ROH_ level was then summed with 0.0625, resulting in an expectation of F_ROH_ = 0.1045 for the offspring of Himba first cousins. Only one individual met this expectation (individual 14 in Figure 4). Although a first cousin preference in arranged marriages has been reported, we did not find any *parental* pairs that were first cousin relatives. Regardless, the presence of long (>5000 kb) ROH segments in many individuals and the distribution of F_ROH_ values resulting from these long ROH segments alone suggests an effect of recent demographic changes.

**Figure 1.**
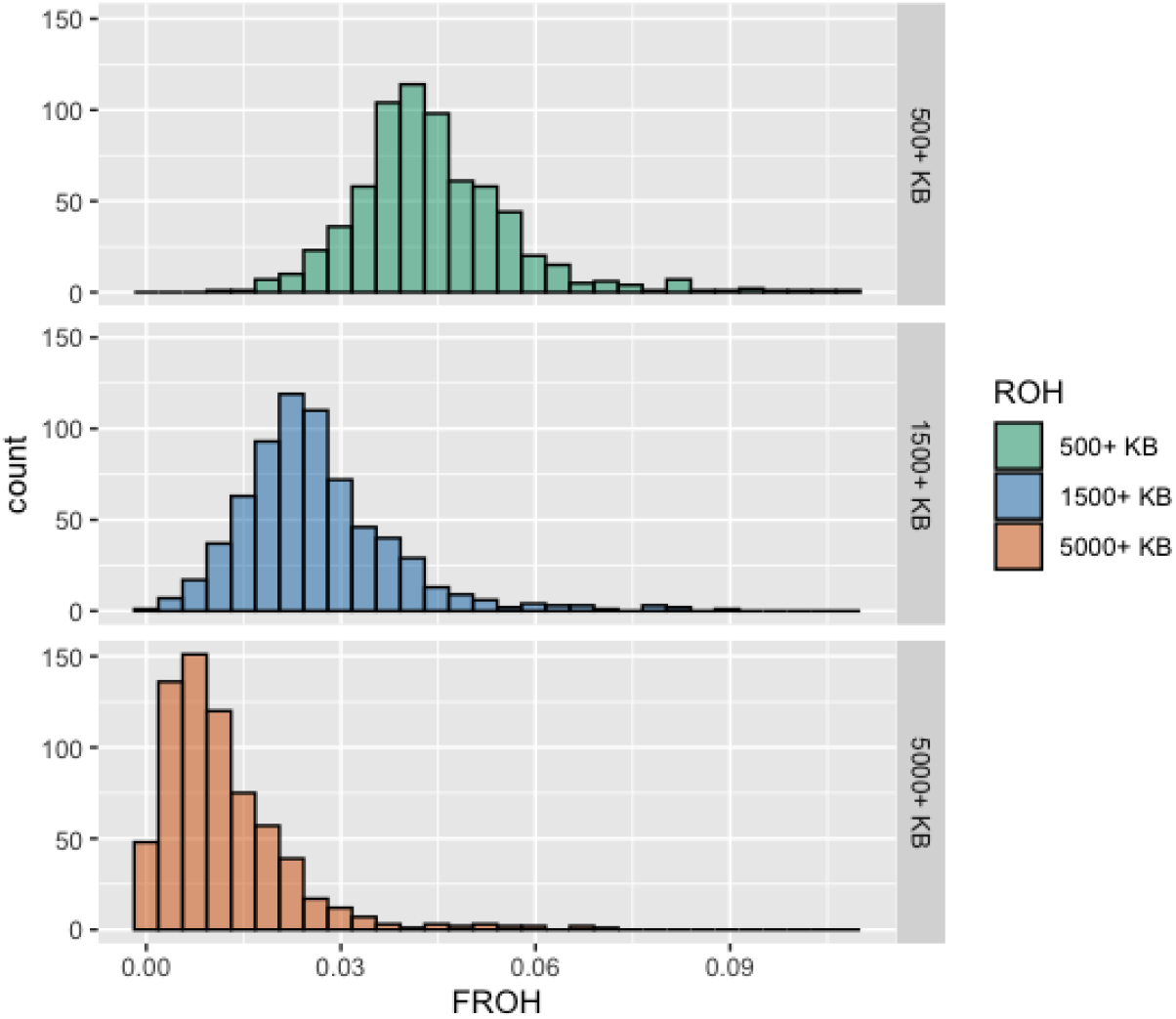
F_ROH_ distributions for varying thresholds to call ROH. The top panel is the distribution resulting when F_ROH_ is calculated using a minimum threshold of 500 kb to call ROH, and the middle and bottom panels are the values calculated using minimum thresholds of 1500 kb and 5000 kb, respectively, to call ROH.

### Inferring a Population Bottleneck

Elevated levels of homozygosity can also result from a population bottleneck. Based on knowledge of historical events, we hypothesized that Himba experienced a recent bottleneck within the past ~6 generations (34). These historical events included heavy cattle raiding beginning in the second half of the 1800s, forcing many Himba out of northern Namibia. In 1897, a severe rinderpest epidemic decimated up to 90% of livestock herds. Contagious Bovine Pleuropneumonia also caused devasting cattle losses in the 1930s and continued to impact herds for at least the next 50 years. Several severe droughts have also occurred throughout the 20^th^ century. Additionally, in the first two decades of the 20^th^ century, the Himba experienced the pressures of genocide, harsh taxes, and decreased mobility and isolation (34). To test this hypothesis of recent bottleneck, we selected 120 unrelated individuals and estimated the effective population size (Ne) for the last 100 generations using a non-parametric method that uses inferred identical-by-descent (IBD) segments between pairs of individuals (35). We optimized the parameters used to infer IBD with a pipeline described in Gopalan et al (36) *(Methods).*

Our estimation of N_e_ through time indicates that a bottleneck has occurred throughout approximately the past 60 generations, reaching a minimum effective population size of approximately 450 individuals 12 generations ago (Figure 2A). Himba exhibit elevated levels of total ROH in the genome compared to other African populations, including representatives from eastern, southern, and western regions: Bench, Zulu, Igbo, and Mandinka. Instead, their distribution is more similar to that of the Chabu foragers from Ethiopia. The Chabu, who are currently experiencing a bottleneck that is estimated to have begun slightly more recently than 60 generations ago (37), have a higher mean value of total ROH, but exhibit a range of values similar to those in the Himba (Figure 2B).

**Figure 2.**
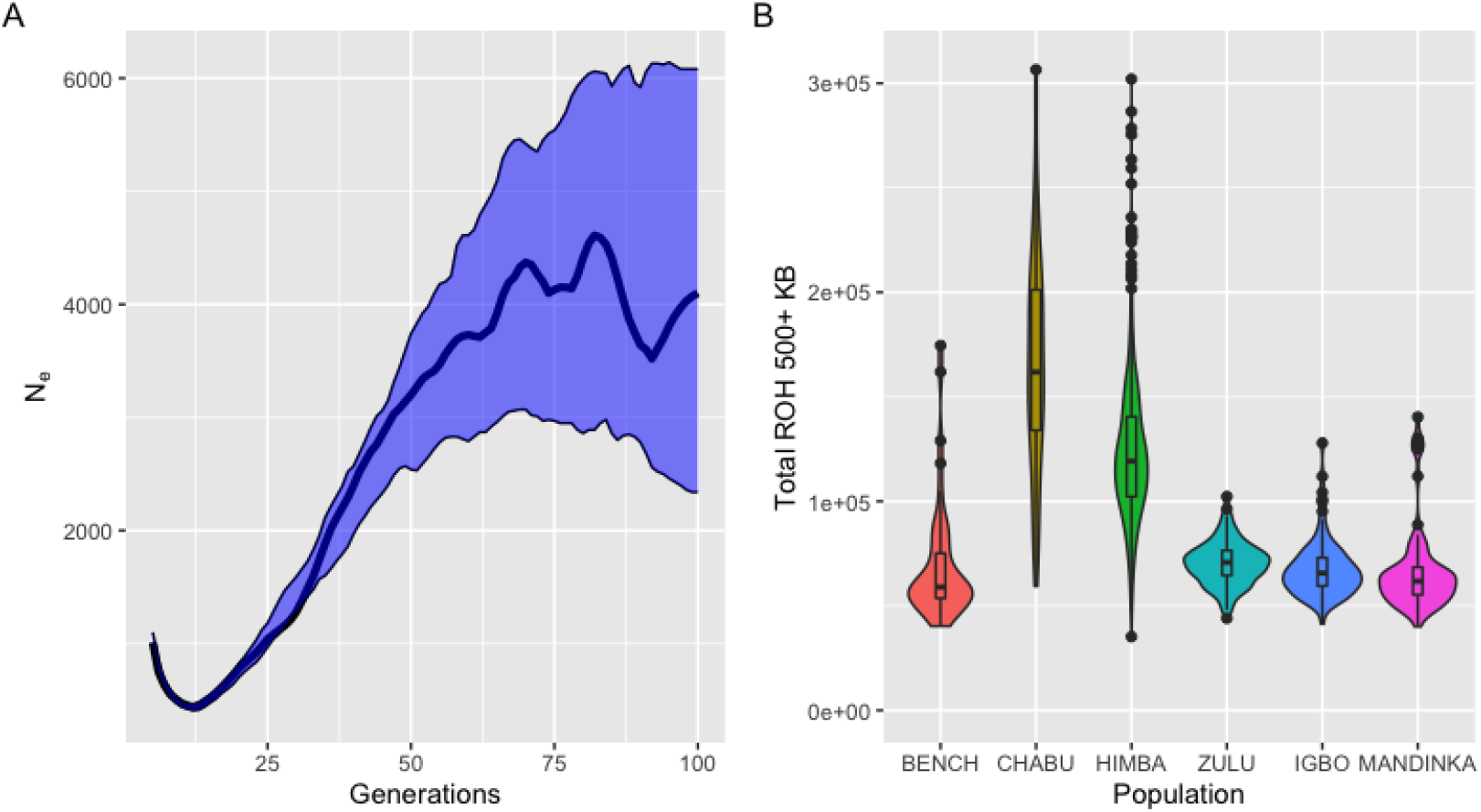
Identical by Descent Segments indicate a recent population bottleneck for the Himba. (a) Effective population size through time was estimated for a sample of unrelated Himba individuals (n=120) using IBDNe. 95% confidence intervals shown in blue. (b) To understand whether the IBD sharing in the Himba was unusual, we contrasted them with two Ethiopian populations (Bench, Chabu), two western African populations (Igbo, Mandenka) and the Bantu-speaking Zulu from South Africa. The distribution of total amount of ROH in individuals’ genomes for ROH segments 500 kb and longer is depicted as a violin plot. The Himba are comparable to the Chabu, an Ethiopian hunter-gatherer population which has also experience recent population decline (Gopalan et al). Strikingly, they have much higher IBD than the Zulu.

To help validate the results of our inferred bottleneck, we simulated two bottlenecks under different scenarios using msprime and compared the IBDNe and F_ROH_ distribution results of the simulations with the actual Himba data. The simulated bottlenecks specified a beginning N_e_ of 4,000 and final N_e_ of 450 to roughly reflect the starting and minimum N_e_ values inferred from the 120 unrelated Himba individuals. In the two scenarios, the starting N_e_ value was set to begin declining at either 60 or 6 generations ago (ga). We find the IBD inferred from the simulated bottleneck beginning 60 ga more closely matches our real data than the one inferred from the simulated bottleneck beginning 6 ga (Figure S1). We also calculated the F_ROH_1500_ distributions for both sets of simulated individuals as well as for our subset of 120 unrelated Himba individuals. We find the F_ROH_ distribution of our 120 unrelated Himba individuals more closely matches the F_ROH_ distribution of our simulated individuals experiencing a bottleneck beginning 60 ga (Figure S2).

### The Effects of F_ROH_ on Fertility

To assess the effects of F_ROH_ on fertility, we identified post-reproductive women in our sample, which we defined as all women greater than 47 years old—as this was the oldest age recorded at which a woman gave birth in this community—and performed a linear regression fit to a Poisson distribution using year of birth (YOB) and number of marriages (NM) as the initial covariates in the modelling process. We used a proxy measure of reproductive success, defined as the number of children who survived to a minimum age of five.

We ran two versions of the model, one including all women regardless of parity, and the other excluding nulliparous women. Many biological conditions, unrelated to recessive load, can affect fertility. In addition to untreated venereal disease (38, 39), conditions such as polycystic ovarian syndrome and endometriosis can affect a woman’s ability to become pregnant (40, 41). When only women who have given birth (n = 65) are considered in the model (i.e. women with zero births are excluded from the model), F_ROH_ has a highly significant (p < 0.006) effect on the total number of children surviving to age five for all three minimum thresholds of ROH. In all three linear models (for each of three ROH thresholds), the number of marriages covariate is eliminated during the backwards elimination and YOB is retained as a significant covariate (Table 1, Figure 3). We find that the effect of F_ROH_ increases with the minimum threshold of ROH, reducing fertility by 10.6%, 12%, and 15%, respectively for F_ROH_500_, F_ROH_1500_, and F_ROH_5000_.

**Table 1.** P-values for variables in models done including fertile women only and including all women indicate a significant effect of F_ROH_ on fertility. The covariates included in each run of the backwards elimination are listed: YOB (year of birth) and NM (number of marriages) and the level of significance is indicated with asterisks.

| Fertile Women (PR women: n=65) |  |  |  |  |  |  |  |
| --- | --- | --- | --- | --- | --- | --- | --- |
| P-value |  | 500+ Kb for ROH |  | 1500+ Kb for ROH |  | 5000+ Kb for ROH |  |
|  | Covariates | YOB | YOB + NM | YOB | YOB + NM | YOB | YOB + NM |
|  | Intercept | 0.02106 * | 0.03676 * | 0.01868 * | 0.03337 * | 0.03094 * | 0.05492 |
|  | F<sub>ROH</sub> | 0.00555 ** | 0.00656 ** | 0.00267 ** | 0.00291 ** | 0.00349 ** | 0.00309 ** |
|  | YOB | 0.01092 * | 0.02163 * | 0.01008 * | 0.02037 * | 0.01803 * | 0.03602 * |
|  | NumMar |  | 0.13360 |  | 0.12189 |  | 0.09272 |
| Effect of F<sub>ROH</sub> |  | 10.6% reduction |  | 12% reduction |  | 15.4% reduction |  |
| All Women (PR women: n=69) |  |  |  |  |  |  |  |
| P-value |  | 500+ Kb for ROH |  | 1500+ Kb for ROH |  | 5000+ Kb for ROH |  |
|  | Covariates | YOB + NM |  | YOB + NM |  | YOB + NM |  |
|  | Intercept | 0.0274 * |  | 0.0264 * |  | 0.0409 * |  |
|  | F<sub>ROH</sub> | 0.0427 * |  | 0.0195 * |  | 0.0134 * |  |
|  | YOB | 0.0171 * |  | 0.0169 * |  | 0.0276 * |  |
|  | NumMar | 0.0211 * |  | 0.0178 * |  | 0.0138 * |  |
| Effect of F<sub>ROH</sub> |  | 7.8% reduction |  | 9.3% reduction |  | 13% reduction |  |

**Figure 3.**
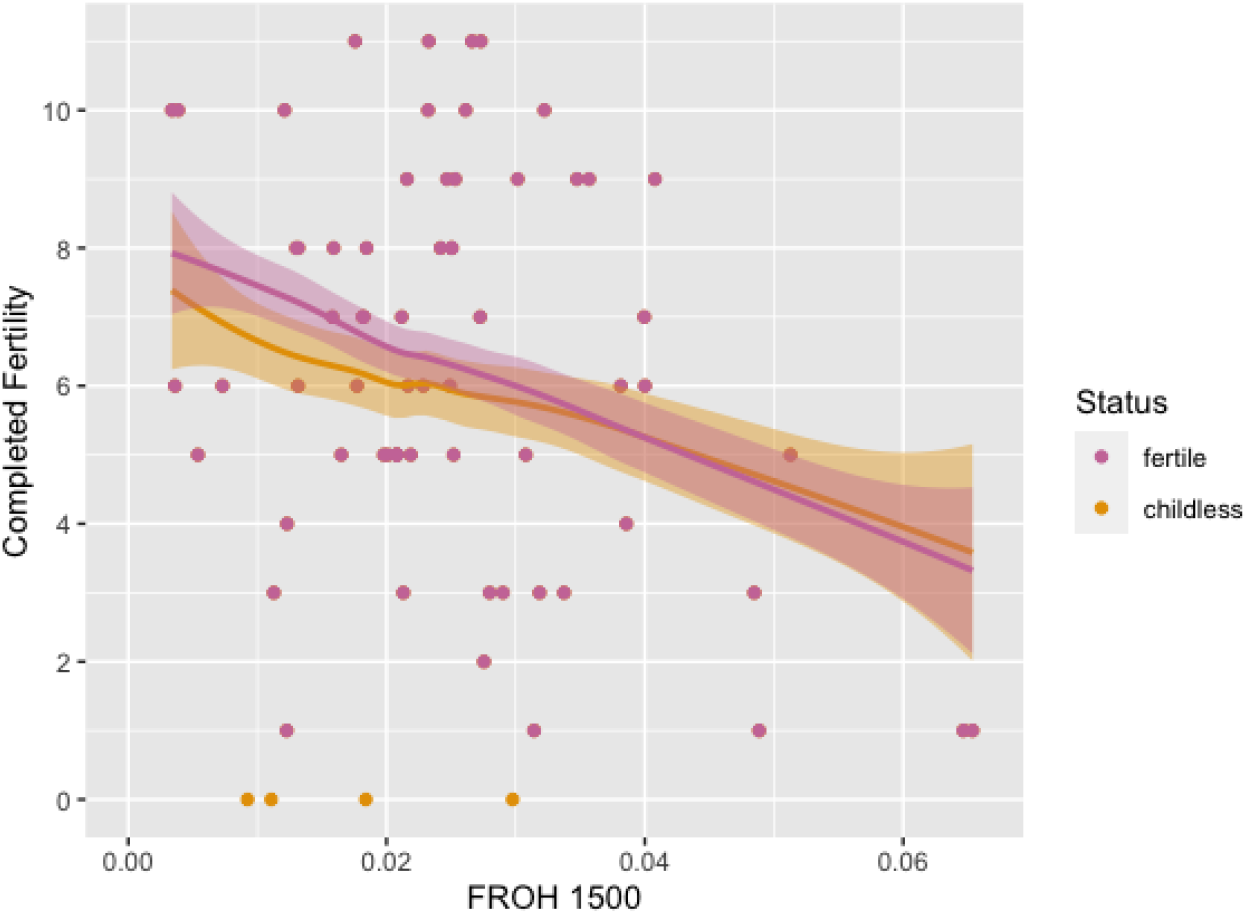
Recent inbreeding is significantly correlated with lower female fertility. The linear relationship between F_ROH_1500_ and completed fertility for post-reproductive women is depicted for both fertile women (i.e. pink line and points) and the additional four women with zero births (i.e. orange points). The fit lines with confidence intervals show the model prediction for the fertile women only (pink) and for all post-reproductive women (orange).

Our sample of all post-reproductive women (n = 69) included four seemingly infertile women (i.e. women who have had zero births). Because we lack medical records that could confirm diagnoses for reduced fertility or sterility, we ran the models again with the addition of the four seemingly sterile women to avoid possible bias. When all women are included in the model, the signal is attenuated, but F_ROH_ continues to have a significant (p < 0.05) effect on completed fertility for all three minimum thresholds of ROH (Table 1, Figure 3). Both covariates had significant effects (p < 0.03), and thus backwards elimination was not performed. We find that the effect of F_ROH_ increases with the minimum threshold of ROH, reducing fertility by 7.8%, 9.3%, and 13%, respectively for F_ROH_500_, F_ROH_1500_, and F_ROH_5000_.

### IBD Sharing between Couples

In order to understand the role of consanguineous marriage on fertility in this population, a closer look at Himba relationship patterns is needed. While cousin marriage is preferred, especially for arranged marriages, “love matches” are also common, particularly following the first marriage. Additionally, it is common and socially acceptable for both husbands and wives to take additional partners (30). Children born outside of marital unions, either through concurrency or out-of-wedlock are referred to as *omoka.* When a woman is married to a man, he becomes the social father of all her children, regardless of biological paternity. Despite this, Himba recognize distinctions between social and biological paternity (42). To assess the effect of different relationships on the distribution of F_ROH_ values, we analyzed the distributions of IBD sharing between different categories of parental couples in a subset of trios (n = 105) for whom we had marriage data, self-reported kinship for married couples, and were able to verify genetic paternity of offspring. Conditional on a couple having confirmed biological offspring together, trios were divided into three categories: A. Married couples self-reported to be unrelated (n = 7), and B. Married couples self-reported to be related *(Methods)* (n = 26). C. Mothers and boyfriends (n = 72). Of these self-reported related couples in “B”, 15 were described as first cousins, 3 were described as first cousins once removed, 1 was described as avuncular, 1 was described as avuncular once removed, and the other 6 were described as more distantly related.

We found no significant difference (p > 0.085) between the distributions of IBD sharing between couples within each category (average IBD between biological parents of *omoka* children = 291 cM, average IBD between unrelated married couples = 347 cM, and average IBD between related married couples = 284 cM). Among married couples purported to be related, the majority of relationships were described as first cousins or first cousins once removed. The average IBD sharing between first cousin pairs (n = 291 pairs) and half-cousin pairs (n = 957) in the Himba, identified using PONDEROSA (32), is 1157 cM and 725 cM, respectively. Thus, while we have many examples of first cousins in the dataset, we observe much lower levels of IBD sharing between married couples. Despite reported relationships, individuals do not appear to marry biologically close kin. However, this does not preclude higher IBD sharing in some couples.

In fact, the parental couples with the highest IBD sharing in our dataset are extra-pair relationships (boyfriend/girlfriend) in which one couple shares a total of 1038 cM (individuals 9 and 11 in Figure 4) and the other shares a total of 1074 cM (individuals 9 and 10 in Figure 4). These couples have IBD consistent with a third-degree relationship but, after reconstructing their pedigree *(Methods),* they are not first-cousins. Rather, they are related through multiple distant relationships. The women (individuals 10 and 11) are paternal half-siblings. They are both half-cousins with the man (individual 9) through their fathers (individuals 4 and 7). Additionally, woman 11 shares a fourth-degree relationship with the man’s paternal grandfather, and the man shares a fourth-degree relationship with the woman 11’s mother. Woman 10’s mother also shares a cryptic relationship with the man’s father (Figure 4). This type of pedigree, where two individuals are connected through multiple distant relationships or “reticulations,” is common within the Himba.

**Figure 4.**
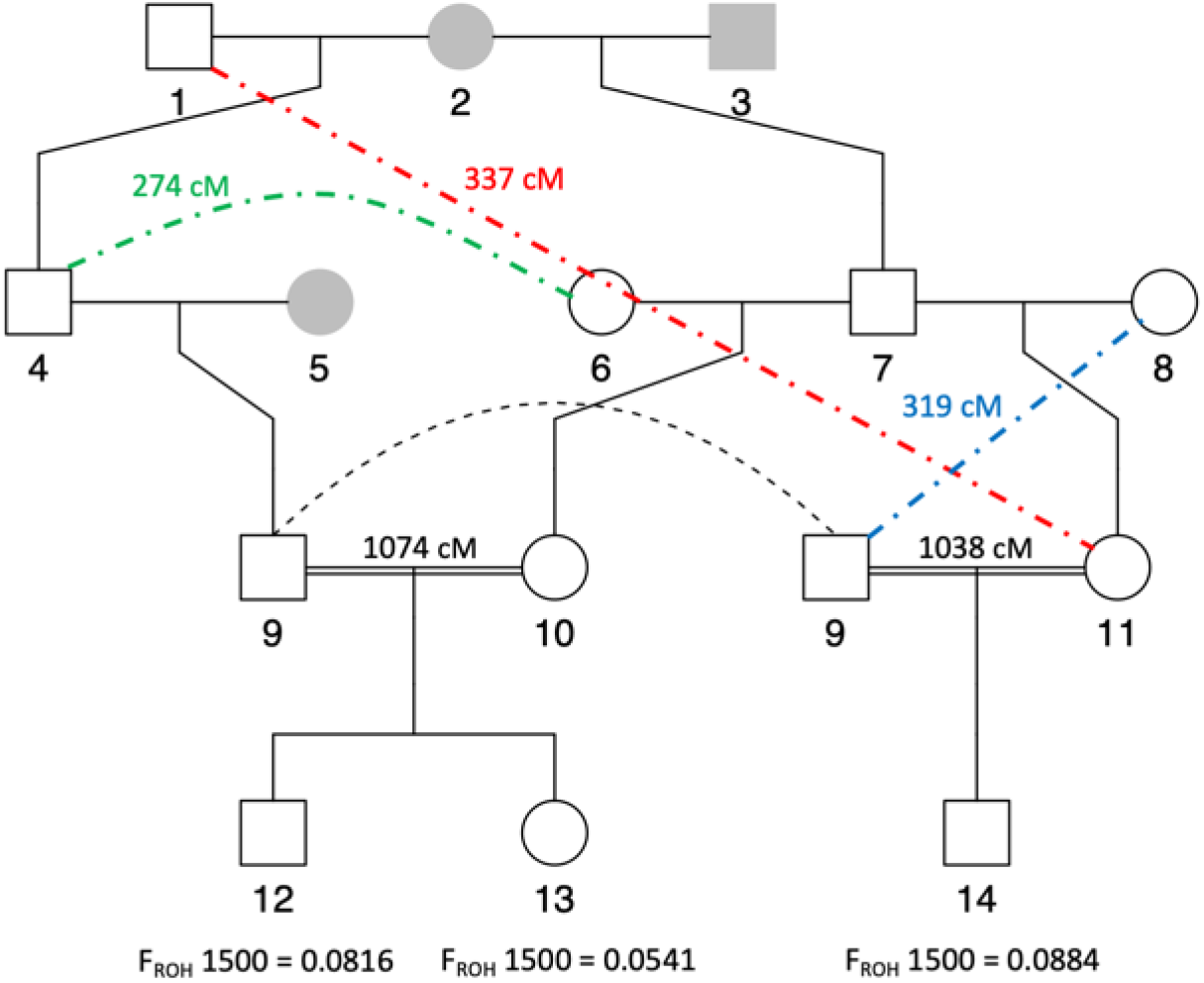
Partial pedigree reconstruction for a couple with high IBD sharing. To parse the relationship between parental couples with high IBD sharing, we reconstructed their pedigrees. Parents 9 and 11, as well as 9 and 10, are half-cousins through their fathers, individuals 4 and 7. Unsampled individuals are shown in gray. The pairwise shared IBD for additional cryptic relationships are represented with colored dashed lines and corresponding values. Additional offspring, siblings, and partners are not shown for the individuals represented here. Individual 9 is represented twice (connected by a black dashed line) for clarity in the representation of the pedigree.

## Discussion

Long runs of homozygosity in human populations can be generated by consanguinity as well as strong founder effects (13, 43–45). Using a minimum threshold of 1500 kb to call ROH, the average fraction of the Himba genome in long runs of homozygosity is 2.6%. Although we observe elevated levels of F_ROH_1500_ and Himba social norms promote consanguineous marriage (46), we do not observe any first cousin parental pairs in our dataset even among married couples who report consanguinity. Rather, our results indicate that the Himba of northern Namibia experienced a population bottleneck that reached its minimum effective population size 12 generations ago. Although historical data suggests a more recent bottleneck (within the past 6 generations), we were unable to detect such a bottleneck here. This may be due to a lack of actual genetic bottlenecking during this time or the difficulty of the IBDNe program to estimate accurate effective population sizes for the most recent generations. The bottleneck we observe reflects a more distant population founding event, which could trace back to the Bantu expansion across southern Africa over the 2,000 years. However, total ROH in the genome is much higher in Himba than in Zulu, suggesting that the Himba elevated ROH is not solely due to the Bantu expansion since the Zulu are also a southern African population derived from the Bantu expansion (47).

Among married couples in our sample, half reported that their partner was a 2^nd^ or 3^rd^ degree relative. However, after reconstructing genetic pedigrees, we observed low levels of IBD sharing among married couples who are purported to be related. There are several possible explanations for the discrepancy between social and biological relatedness among consanguineous couples. First, errors in the demographic interviews could account for some differences, if individuals reported spouses as a first cousin, when in fact they were a more distant relation. However, another likely explanation is that a high rate of extra-pair paternity has led to an untethering of social and biological relatedness. Himba have a strong cultural tradition of sexual concurrency, with most adults having both marital and non-marital partners (48), and previous analyses have shown that Himba have an extra-pair paternity rate of 48% (31). Concurrency is presumed to be a longstanding practice, as it was written about in the first ethnographies of this group in the early 20^th^ century (49–51). This means that if a man marries his cross-cousin (e.g. his mother’s brother’s daughter), there is only about a 50/50 chance she is his biological cousin. With successive generations of concurrency and extra-pair paternity, the chance of relatedness reduces even further. Over time, this could lead to the results we show here. This interplay between concurrency and a preference for consanguinity may thus allow Himba to reap the benefits of a densely connected kin network, while minimizing the costs of higher mutation load that might typically come with consanguineous marriage.

The presence of long ROH is particularly important in assessing the effects of mutation load on fertility. Increased risk for complex diseases has been shown in populations where IBD sharing is elevated or consanguinity is common, suggesting a causal role for multiple recessive mutations throughout the genome (12). Furthermore, long ROH have been shown to be enriched for deleterious nonsynonymous homozygous derived genotypes (17, 43). Therefore, it has been suggested that fitness could be reduced via gene knockouts within long ROH (43). Although our study was limited in its sample size of post-reproductive women, we found F_ROH_ to have a significant effect on completed fertility. Our results show a trend of increasing effect size of FROH on fertility when longer and longer runs of homozygosity are analyzed. This result is consistent with longer tracts carrying greater numbers of deleterious mutations or more damaging mutations such as gene knockouts. Additional variance in phenotypic effect may be caused by varying placement of ROH within the genome. Specific genomic regions may be especially harmful to reproduction if deleterious variants disrupt genes critical to proper reproductive function (52, 53). Overall, our results suggest a multi-locus effect on fitness driven by the expression of deleterious recessive alleles, especially those harbored in long ROH.

Our results are consistent with earlier self-reported pedigrees studies which considered the effect of consanguinity on fertility. Chagnon et al (21) analyzed the South American Yanomamö group and reported that children whose parents were more closely related (i.e. children who would be expected to have higher FROH) had significantly lower fertility themselves. Similarly, Postma et al (22) used town records to reconstruct genealogies for individuals from a small Swiss village, and reported inbreeding depression for fertility among women. Our results strengthen these findings by demonstrating this pattern in another human population using molecular genetic measurements rather than pedigree estimates of relatedness. We caution that self-reported pedigrees may not always reflect biological kinship coefficients, as discussed above.

Variance in fertility within our cohort of post-reproductive women is not solely due to recessive genetic effects; our models explain 8-13% of the variance indicating fertility is affected by other factors. Secondary sterility resulting from untreated venereal disease, such as gonorrhea, may be a factor in this population (38, 39, 54). Pennington and Harpending (38) have suggested that, prior to the 1960s, venereal disease caused lower fertility in the Namibian Herero, a population closely related to the Himba. They argue that women born after ~1945 would be at low risk for sterility due to venereal disease since antibiotics would have been available by the time they became sexually active, and women born prior to ~1915 would be at the greatest risk since antibiotics would not have been available until after they reached menopause (38). In our sample, nine women were born prior to 1945, including one of the four infertile women (YOB = 1943) and none of the women in our sample were born prior to 1915 (minimum YOB = 1926).

However, more recent work by Hazel et al (54) has shown the continued prevalence of gonorrhea infection along with low levels of treatment among Kaokaoland pastoralists, including Himba. Furthermore, this study suggests that these infections may be limiting fertility, but it is unclear to what extent, if any, this is occurring as no official measurements or analyses of the effects of gonorrhea on fertility have been made. A lack of medical records in our Himba sample makes it impossible to know whether there are other exogenous medical conditions that could be affecting fertility as well.

Our results are important for understanding the architecture of complex traits in human populations by suggesting that recessive mutation load may play a key role in fitness. A recent study by Szpiech et al (55) noted that ROHs from different ancestries had different proportions of damaging homozygotes. This is thought to be due to differing population histories, resulting in haplotypes from high heterozygosity populations (such as African populations) containing more strongly deleterious variants than those found on haplotypes from lower-heterozygosity populations, and thus being more severely damaging when found in ROH. The effects of F_ROH_ on fertility and other polygenic traits may then differ between populations. Thus, this work should ideally be replicated in several populations of varying demographic histories. Future work pertaining to the effects of F_ROH_ on fertility should also annotate variants found in ROH to assess their level of deleteriousness. In conclusion, this work is especially important in demonstrating how differences in mutation loads, especially when considered under a recessive model, may affect differences in evolutionary fitness.

## Materials and Methods

### Study Population

The Himba are a small Bantu-speaking population that resides in northwestern Namibia. They are a seminomadic agro-pastoralist group who practice polygyny with a reported first-cousin preference for arranged marriages; however, “love matches” are also common. Additionally, it is common and socially accepted for both husbands and wives to take additional partners (30). Within the past 170 years, the Himba have experienced many factors that have likely contributed to population decline, including genocide, severe drought, and rinderpest epidemics that decimated cattle resources (34).

### Ethical Approval

The data in this study were collected following ethical approval granted by the University of California, Los Angeles (IRB-10-000238) and the State University of New York, Stony Brooke (IRB-636415-12). The study was also approved by the Namibian Ministry of Home Affairs and the University of Namibia Office of Academic Affairs and Research, and local approval of the study was granted by Chief Basekama Ngombe. These data were collected as part of the Kunene Rural Health and Demography Project, which has been working in the community since 2010. Community leaders were actively involved in discussions regarding the research, including who could access the genetic data and what it could be used for, prior to data collection (with initial consultation in 2013 by BMH, BAS; followed by subsequent consultation after DNA collection in 2016). All data and samples were collected with informed consent and parental assent for minors.

### Genetic Data

Individuals were genotyped on either the MEGAex or H3Africa SNP-array. Quality control and filtering proceeded as documented in dbGaP phs001995.v1.p1. We obtained genetic data from three more individuals on the H3Africa array and added them to the dataset after initial QC steps by filtering for SNPs common between the original dataset and the three new individuals. Both datasets were filtered to contain autosomes only to avoid X-chromosome interference in F_ROH_ estimates for males. Genotype data from the H3Africa array was thinned using PLINK2/1.9 to match the SNP density of the MEGAex array, therefore ensuring consistency when calling ROH and allowing for higher SNP-density than would be present after merging the datasets due to missingness across platforms. The final H3Africa dataset contained 504 individuals and 755,660 SNPs and the final MEGAex dataset contained 177 individuals and 755,423 SNPs.

We performed a test to ensure that randomly thinning the SNP density did not affect the estimation of an individual’s F_ROH_. To do this, we thinned the H3Africa dataset 10 separate times, identified ROH in each set as before with a minimum segment length threshold of 1500 kb, and recalculated F_ROH_ in each set. We then calculated the absolute value of the differences between the F_ROH_ values in each new thinned set and in the original thinned set for each individual. We averaged the 10 values for differences between tests for each individual, resulting in an average difference for F_ROH_ calculated from differently thinned SNP array data for each individual and used these values to calculate a root-mean-square error (RMSE) of 0.00058, confirming no significant difference between tests.

### Measures of Inbreeding

To calculate F_ROH_, we first calculated the length of the genome tested in the SNP-arrays by summing the lengths between the first and last SNPs genotyped on each chromosome. We then identified runs of homozygosity (ROH) in each individual using PLINK2/1.9 and calculated F_ROH_ by dividing the total length of the genome found to be in ROH by the total length of the genome tested in the SNP array. F_ROH_ was calculated for each of three different minimum lengths to call ROH: 500 kb, 1500 kb, and 5000 kb. These calculations were done separately for both platforms and then combined in R, resulting in F_ROH_ estimates for a total of 681 Himba individuals.

### Measure of Reproductive Success

Reproductive success was measured as the number of children a woman had who survived to a minimum age of five years old and these data were collected during interviews. In populations where infant and child mortality are high, survival to age five is used as a proxy for likelihood of survival to adulthood. In addition, still births and infant mortality are sensitive topics, which women were often reticent to report. Thus, a measure of fitness based on number of births alone is both less reliable and less pertinent to overall reproductive success than number of children surviving to age five.

### Linear Modelling and Statistical Analysis

To assess the effects of F_ROH_ on fertility, we performed a Poisson linear regression (GLM) in R following a backwards elimination protocol when covariates were insignificant. We assessed the effects of F_ROH_ on fertility for three different measurements of F_ROH_ (using minimum thresholds of 500 kb, 1500 kb, and 5000 kb to call ROH) and used the measure of total number of children surviving to age five as the response variable. Year of birth (YOB) and Number of Marriages (NM) were used as covariates in the modelling process.

### IBDNe

To determine the effective population size through time, we first identified a subset of unrelated individuals (n=120) and extracted them from the larger, already-quality-controlled (see Genetic Data) dataset. We used a custom pipeline (37) to phase, join IBD segments that may have been split due to errors, compare to ROH, and choose the optimal parameters for input into the IBDNe program following a pipeline described in Gopalan et al (2020) (36), which culminates with the running of IBDNe (35). The output was graphed in R.

Our msprime simulations followed Gopalan et al (2022) (37) and specified a starting Ne, a final N_e_ in the present, the generation at which to begin the bottleneck, and the number of haploid individuals to simulate. This changing N_e_ simulation uses a coalescent model with recombination (“Hudson” model) until 100 ga, then switches to a discrete Wright-Fisher model (37). Our starting and final N_e_ values were 4000 and 450, respectively, to reflect the approximate starting and minimum N_e_ values inferred by IBDNe in the real data. We chose to simulate a bottleneck beginning 60 ga to reflect what was inferred from our actual data, as well as a more recent historical bottleneck beginning 6 ga to reflect our original hypothesis. We simulated 120 diploid individuals to match the number of Himba individuals used to infer the bottleneck in our actual data. Because simulated data do not have the same issues as actual data (i.e. they are perfectly phased and without genotype errors or missing SNPs), we did not need to use the same custom pipeline to optimize the parameters to infer IBD and run IBDNe. Instead, we processed the vcf output from the simulations, identified IBD with hap-ibd, and ran IBDNe following Gopalan et al (2022) (37).

### Paternity Identification and Marriage Data

*Omoka* status was determined and confirmed with genetic paternity analysis as described in Scelza et al (31). Children were described as *omoka* if their biological father was not their mother’s husband at the time of their birth. Information regarding marital unions and the descriptions of married couples’ relatedness were collected in interviews and then translated into a specific category of relatedness. For example, a description of “mother’s brother’s daughter” was translated to the category of first cousins, while descriptions such as “father’s brother’s daughter’s daughter” or “the husband’s mother is from the wife’s paternal uncle” were translated to the category of first cousins once removed. One couple was described to be related in two different ways, as both first cousins once removed and possibly as third cousins once removed as well. However, it was unclear to what extent these two connections were independent or overlapping so they were included in the category of the closer of the two relationships (i.e. first cousins once removed).

### Pedigrees

To reconstruct the pedigree for our high IBD couples, we used PONDEROSA, an algorithm that infers pedigree relationships and is especially suited for populations with elevated IBD sharing (32). The Kinship2 R package was used to plot the relationships identified by PONDEROSA.

## Acknowledgements and Funding

This research was funded by the NSF (BCS-1534682 to B.A.S) and NIH (R35GM133531 to B.M.H). We would like to thank the Himba for their participation and hospitality.

## Supplemental Figures

**Figure S1.**
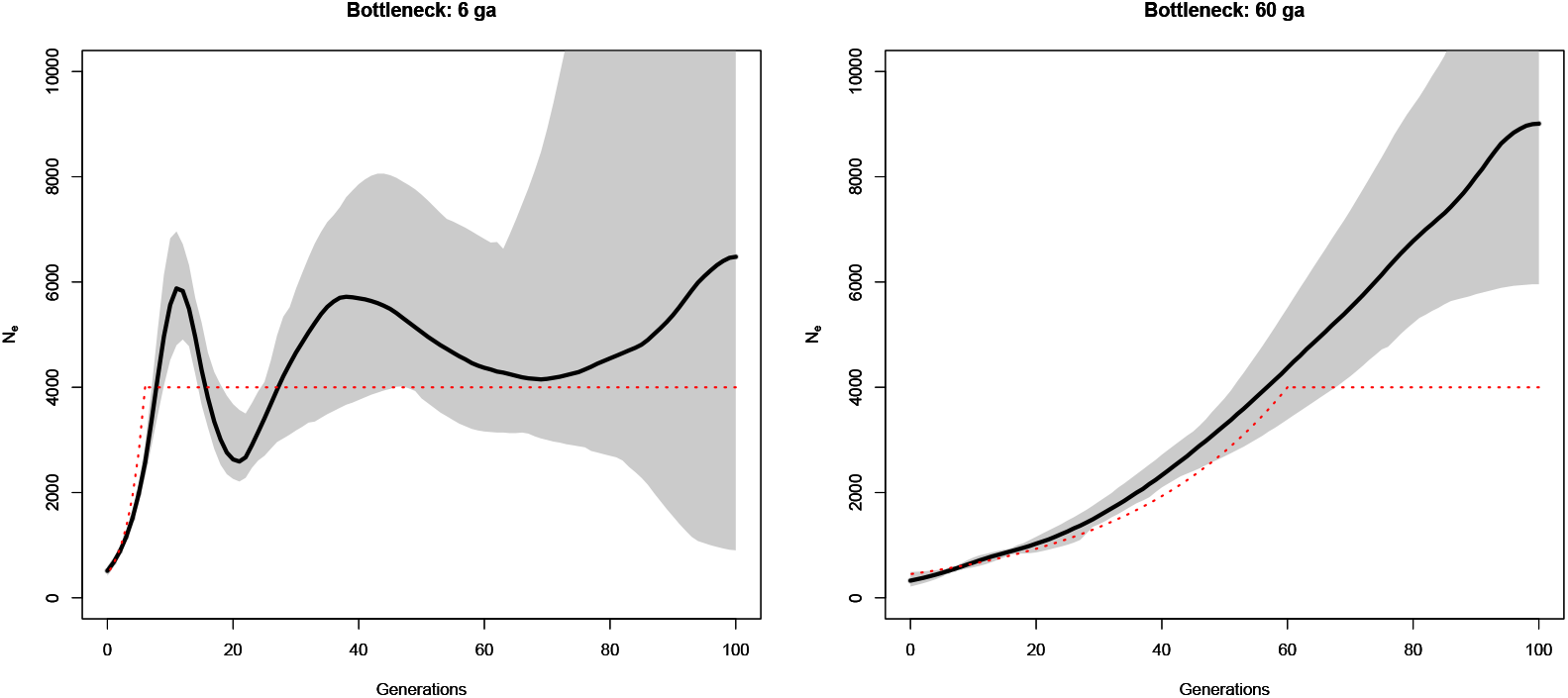
To help validate the results of our inferred bottleneck, we simulated two populations experiencing a bottleneck beginning either 6 ga or 60 ga in msprime and used IBDNe to infer the simulated bottleneck. The black line and 95% confidence intervals (shown in gray) are inferred by IBDNe, and the red dashed lines represent the simulation parameters.

**Figure S2.**
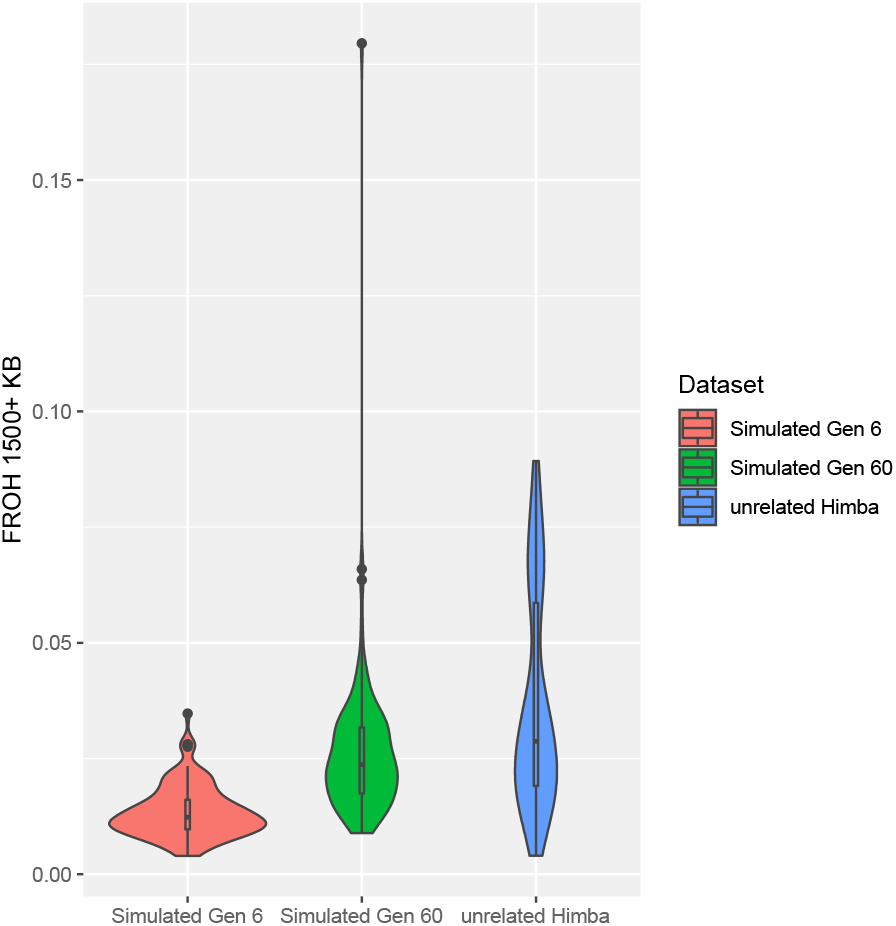
To understand how bottlenecking beginning at different time points produces different distributions of F_ROH_ and validate the timing of our inferred bottleneck, we compared F_ROH_1500_ distributions between simulated and actual individuals. The set of unrelated (n=120) Himba individuals are shown in blue and the simulated individuals with bottlenecks beginning at 6 and 60 generations ago are shown in red and green, respectively.

## Notes

https://www.ncbi.nlm.nih.gov/projects/gap/cgi-bin/study.cgi?study_id=phs001995.v1.p1

### Competing Interest Statement

The authors have declared no competing interest.

